# Lycium barbarum polysaccharide alleviates neurobehavioral deficits in mice with ischemic cerebral injury

**DOI:** 10.1101/2025.06.13.659653

**Authors:** Ruiqing Mian, Liqiong Ma

## Abstract

**Objective:** To investigate the effects of Lycium barbarum polysaccharides (LBP) on neurobehavioral impairments in mice with ischemic stroke and explore the underlying mechanisms.

**Methods:** A middle cerebral artery occlusion (MCAO) model was established using the filament occlusion method to evaluate the therapeutic effects of LBP on pathological brain tissue damage after cerebral ischemia-reperfusion injury (I/R). Mice were randomly divided into three groups: sham surgery (Sham), I/R, and I/R + LBP. Behavioral tests, including the Y-maze test, rotarod test, and balance beam test, were systematically conducted to assess the impact of LBP on neurobehavioral impairments. Enzyme-linked immunosorbent assay (ELISA) was used to quantify peripheral blood levels of pro-inflammatory cytokines tumor necrosis factor-α (TNF-α) and interleukin-6 (IL-6), reflecting inflammatory status.

**Results:** LBP significantly ameliorated neuroinflammation and oxidative stress in mice with cerebral I/R injury, demonstrating marked protection against I/R-induced neurofunctional damage. LBP notably improved motor and memory deficits caused by ischemic stroke. Compared to the I/R group, LBP improved neuroinflammation and oxidative stress levels post-I/R injury.

**Conclusion:** This study demonstrates that LBP alleviates ischemic stroke-induced neurological damage by attenuating inflammatory responses.

## 1. Introduction

Global incidence of stroke continues to rise, imposing a substantial socioeconomic burden due to its high mortality and disability rates [1-3]. Ischemic stroke accounts for the largest proportion, characterized by abrupt onset and a narrow therapeutic time window. Failure to administer standardized treatment promptly may significantly increase the risk of adverse outcomes.

Recent studies have revealed that the core pathological mechanism of ischemic stroke involves: following the rupture of atherosclerotic plaques in large vessels (e.g., carotid artery, middle cerebral artery), local platelet aggregation forms thrombi, leading to a precipitous decline in cerebral blood flow. The subsequent interruption of oxygen and nutrient supply triggers a vicious cycle of intracellular calcium overload-oxidative stress-mitochondrial dysfunction, causing irreversible neurological deficits [4,5]. Therefore, intervention targeting key nodes of the post-ischemic cascade may represent a potential therapeutic strategy to mitigate secondary brain injury.

As a traditional Chinese medicine and dietary supplement, Lycium barbarum has been widely used in disease prevention and health management in China and globally [6]. Lycium barbarum polysaccharides (LBP), a complex polysaccharide fraction extracted from Lycium barbarum fruits (containing 30% polysaccharides and proteins), represent one of the most bioactive components. Existing evidence confirms that LBP exerts multiple pharmacological effects with favorable clinical safety. Leveraging its prominent anti-inflammatory and antioxidant properties, LBP has been applied to ameliorate various pathological processes, including doxorubicin-induced cardiotoxicity, oxidative liver injury, and retinal ischemia-reperfusion (I/R) injury [7-9]. Notably, recent studies have demonstrated that LBP can cross the blood-brain barrier (BBB) to exert neuroprotective effects, providing a theoretical basis for its application in neurological disorders [10].This study aims to investigate the effects of LBP on neurobehavioral deficits in mice with ischemic stroke and explore the underlying mechanisms.

## 2. Materials and methods

### 2.1 Experimental Animals

Twenty-seven adult male C57BL/6J mice (6–8 weeks old) were obtained from the Medical Experimental Animal Center of Shandong Second Medical University. All animals were housed in a barrier-grade animal facility maintained at (22±2) °C with (55±5)% relative humidity under a 12-h light/dark cycle. Mice had free access to food and water throughout the experiment.

### 2.2 Reagents and Instruments

The small animal anesthesia ventilator, Y-maze, and balance beam were purchased from RWD Life Technology Co., Ltd. Isoflurane was obtained from MedChem Express. Mouse IL-6 and TNF-α ELISA kits were purchased from Shanghai Jianglai Biotechnology Co., Ltd. The malondialdehyde (MDA) detection kit and total superoxide dismutase (SOD) activity assay kit were obtained from Beyotime Biotechnology.

### 2.3 Middle Cerebral Artery Occlusion (MCAO) Model Establishment and Drug Treatment

Mice were anesthetized with 1.2% isoflurane, and the right common carotid artery, external carotid artery, and internal carotid artery (ICA) were exposed. A silicone-coated filament (diameter: 0.21±0.02 mm) was inserted into the ICA to occlude the middle cerebral artery (MCA) for 60 min, followed by reperfusion upon filament removal. Sham-operated mice underwent the same procedures without MCA occlusion. LBP was dissolved in PBS, and the MCAO+LBP group received daily gavage administration of 100 mg/kg/day LBP for 14 days prior to surgery.

Behavioral assessments were conducted 24 h after reperfusion. Mice were then deeply anesthetized with 2% isoflurane and euthanized by cervical dislocation. Blood samples were collected via intracardiac puncture, centrifuged at 3000 × g for 15 min at 4°C to separate serum, and stored at -80°C for subsequent analysis.

### 2.4 Modified Neurological Symptom Score (mNSS)

The mNSS system used an 18-point quantitative scale, where 0 points indicated normal neurological function and 18 points represented the most severe deficits. The comprehensive evaluation included four functional domains: 1) motor function assessment (0–9 points), 2) sensory function testing (0–4 points), 3) reflex function evaluation (0–4 points), and 4) balance and coordination testing (0–3 points).

### 2.5 Rotarod Test

Mice underwent strict adaptive training before the test, including stationary rod exploration and low-speed rotation training, followed by formal testing in an accelerating mode. The test was performed 24 h after MCAO, with each mouse tested three times; the mean latency and maximum (tolerance speed) were recorded. The experiment was conducted in compliance with animal ethics guidelines, with environmental noise controlled at <50 dB and temperature/humidity maintained at 22–25°C to minimize external interference.

### 2.6 Balance Beam Test

A circular wooden beam (80 cm length, 10–15 mm diameter) was fixed between adjustable stands at a height of 30–50 cm, with a soft pad placed beneath to prevent fall injury. The beam ended in a dark box as a safety target. For 3 days prior to testing, mice were trained to walk from the start to the dark box 5 times daily to establish behavioral memory. During formal testing, the distance traveled along the beam within 60 s, number of limb slips, and falls were recorded for each mouse (3 trials, mean value used).

### 2.7 ELISA analyses

Fresh brain tissue was harvested, followed by the addition of PBS to extract the supernatant for the measurement of TNF-α, IL-6 in the brain tissue were detected via ELISA kits (Jianglai Biotechnology Company, Shanghai, China).

### 2.8 Statistical analyses

Statistical analyses were performed using GraphPad Prism version 9.0. To evaluate differences among the groups, independent two-sample t-tests and one-way ANOVA were utilized. In instances of significant ANOVA results, pairwise comparisons were carried out using Tukey’s post-hoc test. A value of *P<0.05, **P<0.01, and ***P<0.001 was established, all data reported as means±SDS.

## 3. Results

### 3.1 LBP Improves Neurological Function Scores in Mice with Ischemic Stroke

Neurological function scores were assessed 24 h after MCAO, revealing that scores in the ischemia/reperfusion (I/R) group were significantly higher than those in the sham operation group, indicating substantial impairment in motor function, sensory function, reflex function, and balance-coordination ability. Notably, mice treated with LBP exhibited significantly lower neurological function scores compared to untreated mice after MCAO (Fig. 1), suggesting that LBP treatment alleviates I/R-induced neuronal injury.

**Fig. 1.**
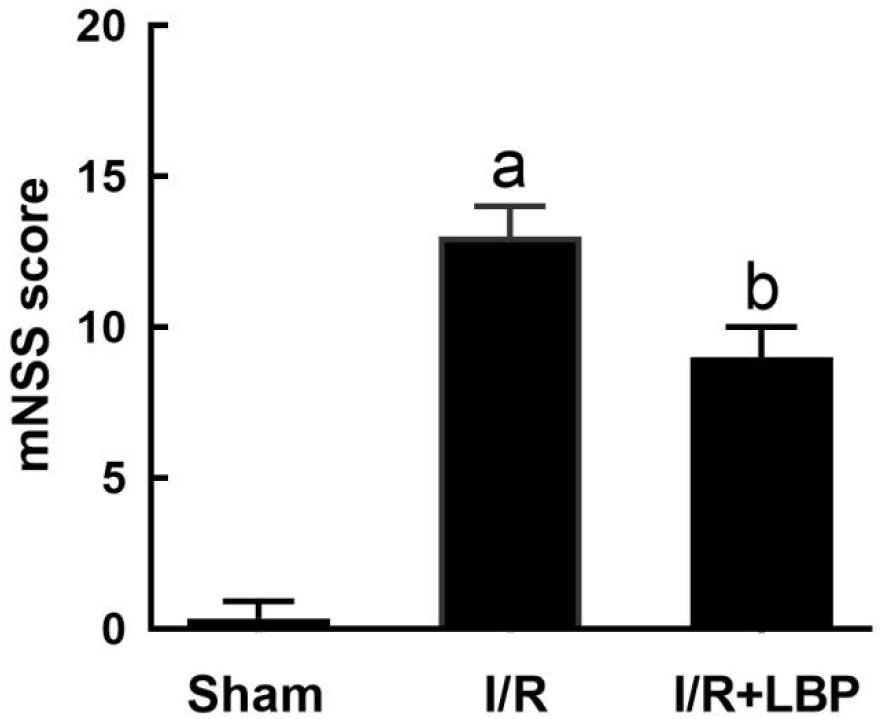
mNSS score of mice. ^a^P < 0.05 vs Sham group, ^b^P < 0.05 vs I/R group.

### 3.2. LBP Improves Learning and Memory Behaviors in Mice with Ischemic Stroke

The Y-maze test is a widely used experimental approach in neuroscience and behavioral research, employed to evaluate spatial working memory, spontaneous exploratory behavior, and cognitive flexibility in rodents. Our study examined Y-maze behavior 24 h after MCAO and found that compared to sham-operated mice, the ischemia/reperfusion (I/R) group exhibited a significant reduction in the number of novel arm entries, indicating spatial memory deficits and decreased exploratory motivation induced by cerebral ischemic injury (Fig. 2). Notably, LBP intervention significantly restored the time spent in the novel arm. These results demonstrate that LBP effectively ameliorates MCAO-induced cognitive dysfunction.

**Fig. 2.**
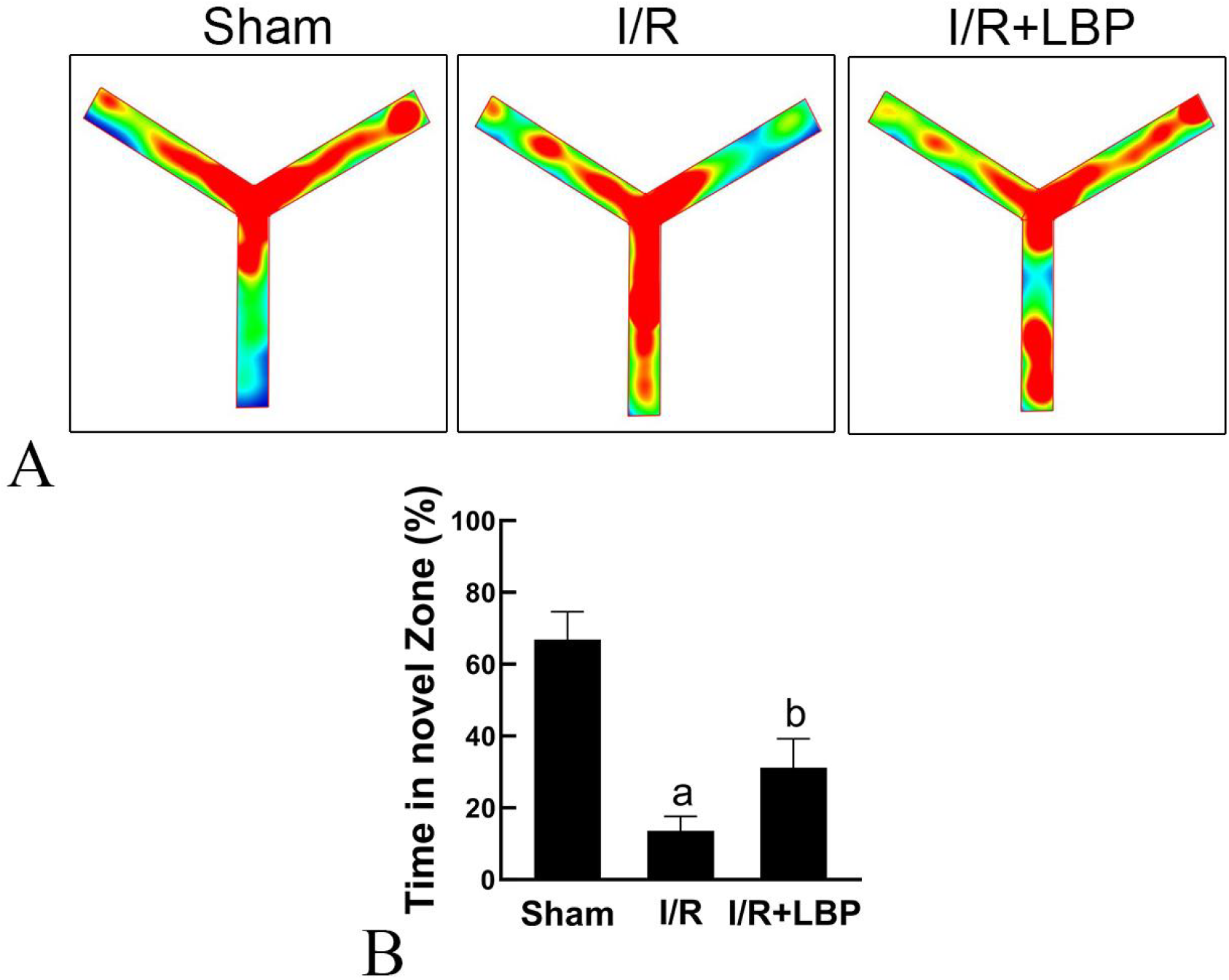
LBP Attenuates MCAO-Induced Learning Impairment. A: Y-Maze Test. B: Time Spent in the Novel Arm. aP < 0.05 vs Sham group, ^b^P < 0.05 vs I/R group.

### 3.3. LBP Improves Motor Behavior in Mice with Ischemic Stroke

To systematically evaluate the effect of LBP on motor function in ischemic mice, motor ability was assessed using the rotarod test and balance beam test. Results showed that compared to sham-operated mice, the ischemia/reperfusion (I/R) group exhibited significantly shorter latency on the rotarod and a significantly reduced total distance walked on the balance beam. Notably, these parameters were significantly improved in LBP-treated mice, indicating that LBP effectively restores motor coordination and balance maintenance after cerebral ischemia (Fig. 3). These behavioral findings further support that LBP exerts neuroprotective effects against ischemic brain injury by mitigating neuronal damage and promoting neuroplasticity.

**Fig. 3.**
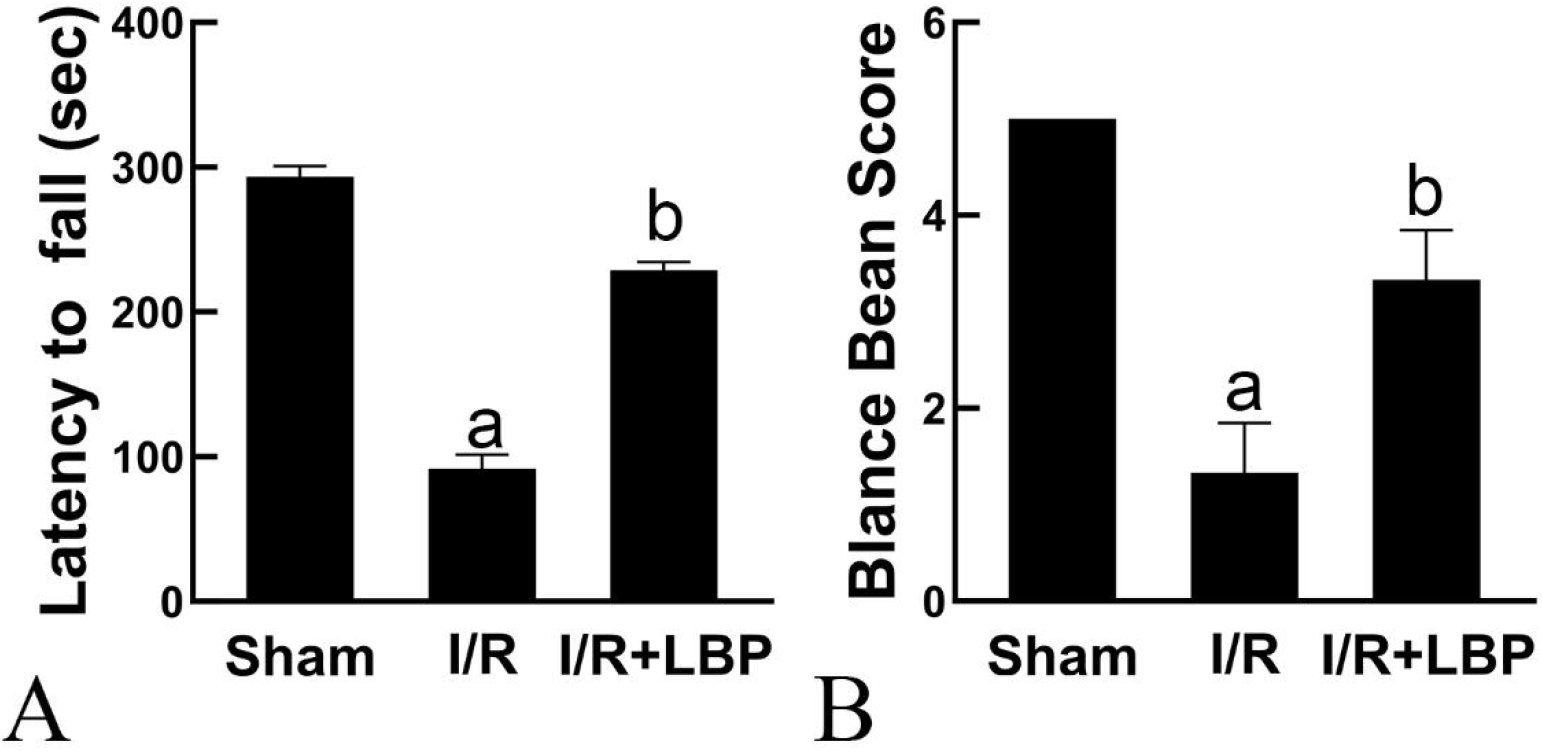
Neurological function score of mice. A: Rotarod test. B: Beam balance test. ^a^P < 0.05 vs Sham group, ^b^P < 0.05 vs I/R group.

### 3.4 LBP Reduces Inflammatory Cytokine Levels in Mice with Ischemic Stroke

In this study, plasma inflammatory cytokine levels were measured by enzyme-linked immunosorbent assay (ELISA) 24 hours post-surgery. Results showed that compared with the sham group, plasma TNF-α and IL-6 concentrations were significantly elevated in the ischemia/reperfusion (I/R) group, indicating that cerebral ischemia triggered a systemic inflammatory cascade. Notably, TNF-α and IL-6 levels in the LBP-treated group were significantly reduced, showing a marked difference from the I/R group (Fig. 4). These findings demonstrate that LBP exerts neuroprotective effects by significantly attenuating neuroinflammation.

**Fig. 4.**
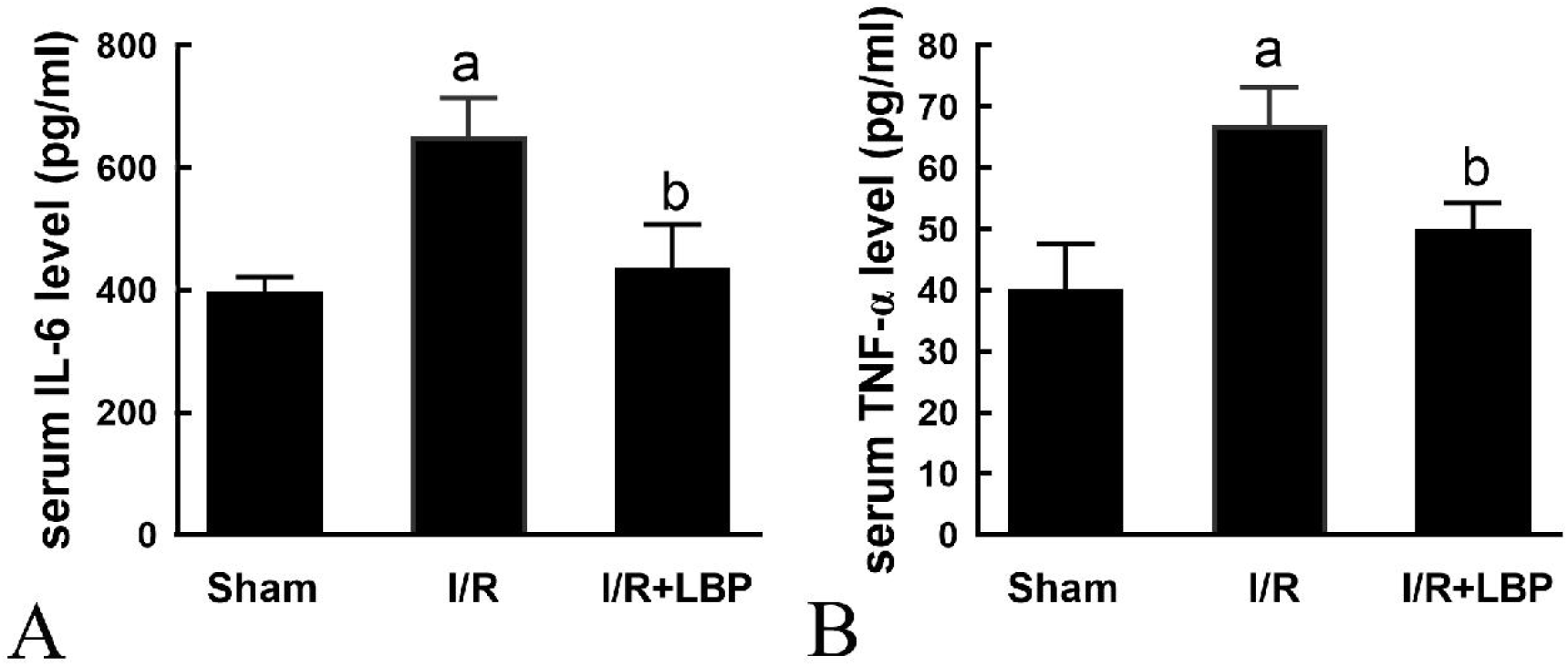
LBP alleviates I/R-induced neuroinflammation. A: Plasma IL-6 Levels. B: Plasma TNF-α Levels. ^a^P < 0.05 vs Sham group, ^b^P < 0.05 vs I/R group.

## 4. Discussion

Stroke, an acute cerebrovascular event, triggers complex pathological cascades following ischemic or hemorrhagic injury, representing the core mechanism underlying progressive neurological deterioration. In ischemic stroke, neuronal energy metabolism failure triggered by blood flow interruption is a key inducer of brain edema formation [11]. Concomitantly, exacerbation of reactive oxygen species (ROS) burst and oxidative stress activates membrane lipid peroxidation, thereby disrupting the blood-brain barrier (BBB) and inducing aberrant release of inflammatory cytokines (e.g., IL-6, TNF-α), which collectively exacerbate neuroinflammation [12]. Without timely intervention, these pathological changes lead to irreversible motor dysfunction, cognitive deficits, and even life-threatening outcomes [13,14]. Thus, developing therapeutic strategies that early intervene and effectively mitigate post-ischemic oxidative stress and neuroinflammation holds critical clinical significance.

Using a middle cerebral artery occlusion (MCAO) model, this study found that ischemic injury not only induced brain edema but also significantly elevated oxidative stress and inflammatory responses in brain tissues. The superimposition of these pathological factors caused remarkable impairments in spatial motor ability and novel object recognition in model mice. Notably, behavioral assays showed that LBP intervention significantly improved motor coordination and spatial memory, providing direct evidence for its neuroprotective effects.

Leveraging LBP’s established anti-inflammatory, antioxidant properties, and excellent BBB permeability, we systematically explored its neuroprotective mechanisms [15-17]. Experimental data showed that LBP intervention effectively alleviated I/R-induced brain edema and improved oxidative stress status, manifested as increased superoxide dismutase (SOD) activity and reduced malondialdehyde (MDA) content—findings suggesting that LBP exerts antioxidant effects by inhibiting excessive ROS production. Importantly, biochemical assays revealed that LBP significantly downregulated key proteins of the NF-κB signaling pathway (IL-6, TNF-α), representing a pivotal molecular mechanism for its coordinated regulation of oxidative stress and inflammation [18-22].

Collectively, these results demonstrate that LBP, as a multi-target neuroprotectant, effectively mitigates neurobehavioral deficits in ischemic stroke by interrupting the “oxidative stress-inflammation” vicious cycle. This provides critical experimental evidence for developing natural product-based stroke therapies.

Notably, this study has limitations: the focus on ischemic stroke models necessitates exploration in diverse disease models. Additionally, LBP’s precise targets and therapeutic time window require further clarification. Future studies should integrate single-cell sequencing and metabolomics to elucidate LBP’s multi-target mechanisms, develop nano-delivery systems to enhance targeting, and explore synergistic effects with clinical thrombolytics—thereby providing robust theoretical support for clinical translation.

## Abbreviations

LB: Lycium barbarum
I/R: ischemia/reperfusion injury
MCAO: transient middle cerebral artery occlusion
ICA: internal carotid artery
MCA: middle cerebral artery
mNss: modified neurological severity score
OFT: open field test

## Declarations

### Ethics approval and consent to participate

All animal experiments in this study were reviewed and approved by the Animal Ethics Committee of Ningxia Medical University.

